# Homogenization of Rhizosphere Bacterial Communities by Pea (*Pisum sativum* L.) Cultivated under Different Conservation Agricultural Practices in the Eastern Himalayas

**DOI:** 10.1101/662775

**Authors:** Diptaraj Chaudhari, Krishnappa Rangappa, Anup Das, Jayanta Layek, Savita Basavaraju, Yogesh Shouche, Praveen Rahi

## Abstract

Conservation agriculture offers a suitable system to harmonize agriculture with the environment, especially in fragile ecosystems of North-East India. Soil microbes play pivotal roles in ecosystem functioning and act as indispensable indicators of overall fitness of crop plant and soil health. Here we demonstrated that altercations in residue management and tillage practices lead to the development of differential bacterial communities forcing the pea plants to recruit special groups of bacteria leading to highly homogenous rhizosphere communities. Pea rhizosphere and bulk soil samples were collected, and bacterial community structure was estimated by 16S rRNA gene amplicon sequencing and predictive functional analysis was performed using Tax4Fun. The effect on pea plants was evident in the bacterial communities as the overall diversity of rhizosphere samples was significantly higher to that of bulk soil samples. *Bacillus, Staphylococcus, Planomicrobium, Enterobacter, Arthrobacter, Nitrobacter, Geobacter*, and *Sphingomonas* were noticed as the most abundant genera in the rhizosphere and bulk soil samples. The abundance of Firmicutes and Proteobacteria altered significantly in the rhizosphere and bulk samples, which was further validated by qPCR. Selection of specific taxa by pea plant was indicated by the higher values of mean proportion of *Rhizobium, Pseudomonas, Pantoea, Nitrobacter, Enterobacter* and *Sphingomonas* in rhizosphere samples, and *Massilia, Paenibacillus* and *Planomicrobium* in bulk soil samples. Tillage and residue management treatments did not significantly alter the bacterial diversity, while their influence was observed on the abundance of few genera. Recorded results revealed that pea plant selects specific taxa into its rhizosphere plausibly to meet its requirements for nutrient uptake and stress amelioration under the different tillage and residue management practices.

## 1. Introduction

Intensive agriculture practiced to meet the food production needs of the ever-increasing population in the changing climate is posing a severe strain to soil and environmental health. Adopting climate-friendly practices appear to be the best strategy to achieve sustainability in agriculture (Rahi, 2017). Conservation agriculture (CA) being an evolved agro-ecosystem management approach for preserving and enriching the environmental resources while improving and sustaining the crop productivity amidst possible environmental stresses (Godfray and Garnett, 2014; Nelson et al., 2017; Muzangwa et al., 2017). Adoption of reduced-tillage or no-till practices preserve soil structure and minimize erosion and loss of organic matter (Hogenhout and Loria, 2008; Mus et al., 2016). Comprehensively, CA requires the simultaneous application of viable crop rotations, minimal soil disturbances, and crop residue retention (Nichols et al., 2015).

North-East India is known for its diverse, risk-prone, and fragile hills and degraded plateau ecosystems. The region is characterized by fragility, marginality, inaccessibility, and ecosystem diversity. Around 77% of the region’s area is hilly and eroded plateaus, with only 12% area under net cultivation. No-till or minimal tillage systems of CA are the alternatives to reconcile agriculture with its environment in these fragile and degraded ecosystems (Ghosh et al., 2010). Incorporation of crop rotation and diversifying cropping systems with pulse crops, including pea, lentil, and chickpea is also an essential step towards CA, which increase the crop yield and decrease the impact of intensive cultivation on the environment (Laik et al., 2014; Gan et al., 2015). Legume crop like peas can enhance soil organic nitrogen contributed by the symbiotic association of legume and rhizobia (Herridge et al., 2008; Rahi et al., 2012; Rahman et al., 2014).

Soil microbial diversity plays a significant role in sustaining ecosystem functioning, as microbial communities influence several ecosystem processes and services. Microbial communities are highly sensitive to soil management practices, and a detailed microbial community analysis can reveal various important indicators of soil quality (Fierer et al., 2007; Sánchez-Cañizares et al., 2017). Previous studies have suggested CA practices like no-till alter microbial diversity and activity significantly when compared to conventional tillage (Degrune et al., 2017; Babin et al., 2019). Similarly, nutrient and residue management practices like the application of chemical fertilizers often result in a change in soil pH and organic carbon content influencing the endogenous microbial communities (Fierer et al., 2007). The rainfed agricultural lands in North-East India is highly susceptible to soil erosion leading poor soil quality. Several strategies including, no-till, minimal tillage and conventional tillage, in-situ residue retention, weed biomass, green leaf manure, and farmyard manure, and rotation of peas and paddy are being tested on experimental farms of the ICAR Research Complex for North Eastern Hill Region, Umiam, Meghalaya, India (Das et al., 2014). Significant increase in soil organic carbon, soil microbial biomass carbon, and dehydrogenase activity has been observed in no-till treatments in comparison to conventional tillage in hills of North Eastern India (Das et al., 2014). Previous studies on CA in North Eastern regions of India suggested that reduction in tillage intensity can reverse soil degradation and improve soil quality, crop productivity and economic returns, but little is known about its impact on the soil microbial community structure.

Crop plants are the other most crucial factor influencing the microbial communities in a CA system (Maarastawi et al., 2018). Plant root exudates stimulate the microbial activity in the rhizosphere, resulting in increased microbial active biomass and abundance in the rhizosphere by several folds in comparison to surrounding bulk soil (Lugtenberg & Kamilova, 2009). During this process of increased microbial activity in plant rhizosphere, some specific microorganisms were also selected, leading to the build-up of plant-specific community in the rhizosphere (Hu et al., 2018). These rhizosphere microorganisms perform various functions to support plant growth and productivity by enhancing nutrient uptake and tolerance against stress, regulating the immune system, and protecting against pathogens (Lundberg et al., 2012). It is expected that crop plants select a specific community of bacteria in their rhizosphere to meet its nutrition and stress requirements, which lead to homogenize the diverse communities developed in response to the selection imposed by various agricultural practices. Based on this we hypothesize that crop plants (i.e., pea) could be the most dominating selection factor in a long-term experiment with multiple tillage and residue management practices in the acidic and iron-rich soils of Meghalaya in the Eastern Himalayas.

## 2. Materials and Methods

### 2.1. Sample collection

Different tillage and residue management treatments are maintained for the last eight years by alternatively growing rice and pea in rotations in the experimental fields of the ICAR Research Complex for NEH Region, Umiam, Meghalaya. Three different tillage treatments, i.e. zero tillage; minimum tillage and conventional tillage were practiced during the cultivation of rice and only zero tillage during pea cultivation. Six treatments of residue management including 100% NPK, 50% NPK, 50% NPK with in-situ residue retention (ISRR), 50% NPK with weed biomass (WB), 50% NPK with green leaf manure (GLM) and 100% organic treatment i.e. farmyard manure (@5t/ha), weed Biomass and rock phosphate (@150 kg/ha) were maintained during rice cultivation. After harvesting rice, seeds of pea was sown at the seed rate of ~80 kg per hectare by opening narrow troughs of optimum depth with the help of manually operated furrow opener in between two rice lines thus giving a row to row spacing of 20 cm for pea. The recommended dose of nutrients and seeds were placed in furrow and covered with soil: FYM mixture (2:1 ratio) for better seed and soil contact. As a whole, only two treatments were maintained for the pea crop, 100 % organic treatment was maintained in the same plots while remaining treatments were replaced with a single inorganic treatment (20:60:40 N:P2O5: K2O kg/ha). Bulk soil and Pea rhizosphere samples were collected from each treatment plot. Bulk soil samples were collected in triplicates and pooled together from all the treatment plots. Three pea plants growing in the mid-rows of each plot were uprooted using a shovel. The whole root systems of the plants were separated from the loosely adhered soil by gentle shaking. The samples from each plot were pooled together in a sterilized polythene bag and transported to the laboratory under 4 °C.

### 2.2. Isolation of rhizosphere soil, DNA extraction, and next-generation sequencing

Roots of pea plants were transferred to the 15 ml sterilized centrifuge tube and submerged in the 10 ml phosphate buffer saline (PBS) to harvest rhizosphere soils. The tubes were subjected to sonication for 60 seconds, and roots were transferred to a fresh centrifuge tube filled with 10 ml PBS. Sonication step was repeated once again to remove the soil adhered to the roots. The process of washing in PBS and sonication was repeated for a total of three times to get the total rhizosphere soil into the PBS. The rhizosphere soil containing PBS was centrifuged at 5,000 x g for 10 min. The soil pellet was used for the extraction of rhizosphere metagenomic DNA. Total community DNA was extracted from bulk soil and rhizosphere soil samples using DNeasy Power Soil kit (Qiagen-Hilden, Germany) following the manufacturer’s instruction. The concentration of resulting DNA was measured using Nanodrop-1000, (Thermo Scientific, USA) and DNA concentration was normalized to 10□ng/μl. The freshly extracted DNA was used as the template for the amplification of the V4 region of bacterial 16S rRNA gene using universal bacterial primers (Fadrosh et al., 2014). The amplicon sequencing (library preparation and sequencing) was performed on the Illumina Miseq platform according to the manufacturer’s instructions (Illumina, USA).

### 2.3. Bioinformatics and statistical analysis

Assembly of forward and reverse reads generated for each sample was carried out using FLASH (Fast Length Adjustment of SHort reads) (Magoc and Salzberg, 2011). Bacterial diversity analysis was done using standard QIIME (v1.9.0) pipeline (Zhernakova et al., 2016) on the assembled sequences. These sequence reads were clustered into operational taxonomic units (OTUs) using UCLUST algorithm (Edgar, 2010) and SILVA database (Quast et al., 2013) by closed reference based OTU picking method keeping 97% sequence similarity threshold. Representative sequences (repset) from each OTU were selected for taxonomic assignment. Statistical analysis of the alpha diversity across different groups and was done using STAMP (Parks et al., 2010). The differential abundance analysis of bacterial genera across the different study groups was also done using STAMP. The abundance of bacterial phylum and genus was represented using GraphPad Prism (GraphPad Software, La Jolla California USA). Beta diversity analysis of the bacterial diversity present in the bulk and rhizosphere soil samples was done using the online tool microbiome analyst (Dhariwal et al., 2017). R language based package ggtern an extension of package ggplot2 was used to plot ternary diagrams of the differential abundant OTUs in the bulk soil and rhizosphere soil samples. Presence of the shared and unique bacterial genera across the bulk soil and rhizosphere samples was investigated using online tool InteractiVenn (http://www.interactivenn.net). OTUs based predictive functional analysis of the bacterial community was done using Tax4Fun package (Aßhauer et al., 2015) in R and KEGG database. Also, Principal component analysis (PCA) based on the abundant predicted functions was performed in the PAST3 (PAleontological STatistics) software (Øyvind et al., 2001).

### 2.4. Absolute quantification of bulk soil and Pea rhizosphere bacteria

Quantitation of total bacteria and phylum Firmicutes was done using respective qPCR primers (**Table 1**). Briefly, for each gene 10 μl reactions (in triplicate) were set containing suitable pair of primers (0.5μM), 10□ng of metagenomic DNA and SYBR green master mix (Applied Biosystems Inc. USA). The reactions were run on 7300 Real-time PCR system (Applied Biosystems Inc. USA) using the following PCR conditions: initial denaturation at 95□°C for 10□min, followed by 40 cycles at 95□°C for 10□s, 60□°C for 1□min. Group-specific standard curves were generated from serial dilutions of a known concentration of individual PCR products. The amplification specificity of primers was checked by melt curve analysis performed at the end of qPCR cycles. Mean values of the three replicate were used for enumerations of tested gene copy numbers for each sample using standard curves.

**Table 1:**
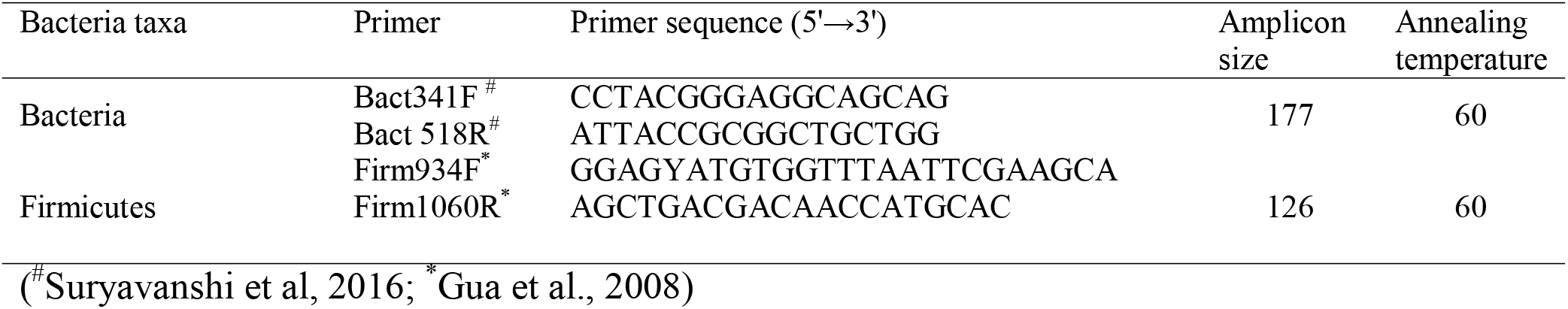
qPCR primers used in this study

### 2.5. Data availability

The sequence data is made available at NCBI SRA submission with accession number SUB5624752 (Bioproject ID: PRJNA544901).

## 3. Results

A total of 2,686,925 raw sequence reads were generated for the bulk soil (n=18) and rhizosphere soil (n=18) samples using the Illumina Miseq sequencing platform. Amongst these sequences, 2,650,426 (98.6%) sequences were assembled using FLASH and clustered into 15,048 OTUs. Among these, 10,078 OTUs were represented by two or more than two sequences, while remaining OTUs were singletons.

### 3.1. Diversity analyses of Bacterial Communities

Alpha diversity assessed by Observed OTUs, Shannon, Simpson, and Chao1 indices differed significantly between rhizosphere and bulk soils (p>0.05) (**Fig. 1**). The rhizosphere exhibited a strong influence on bacterial diversity across various treatments, as the values of diversity indices were always significantly higher in rhizosphere samples in comparison to bulk soils (**Fig. 1**). The factors tillage and residue management practices do not influence alpha-diversity indices (**Fig. 1**). Both bulk soil and rhizosphere soil samples from the organic treatment exhibited least inter-sample variation in Shannon and Simpson alpha diversity indices, in comparison to other residue management treatments (**Fig. 1h and1k**).

**Fig. 1.**
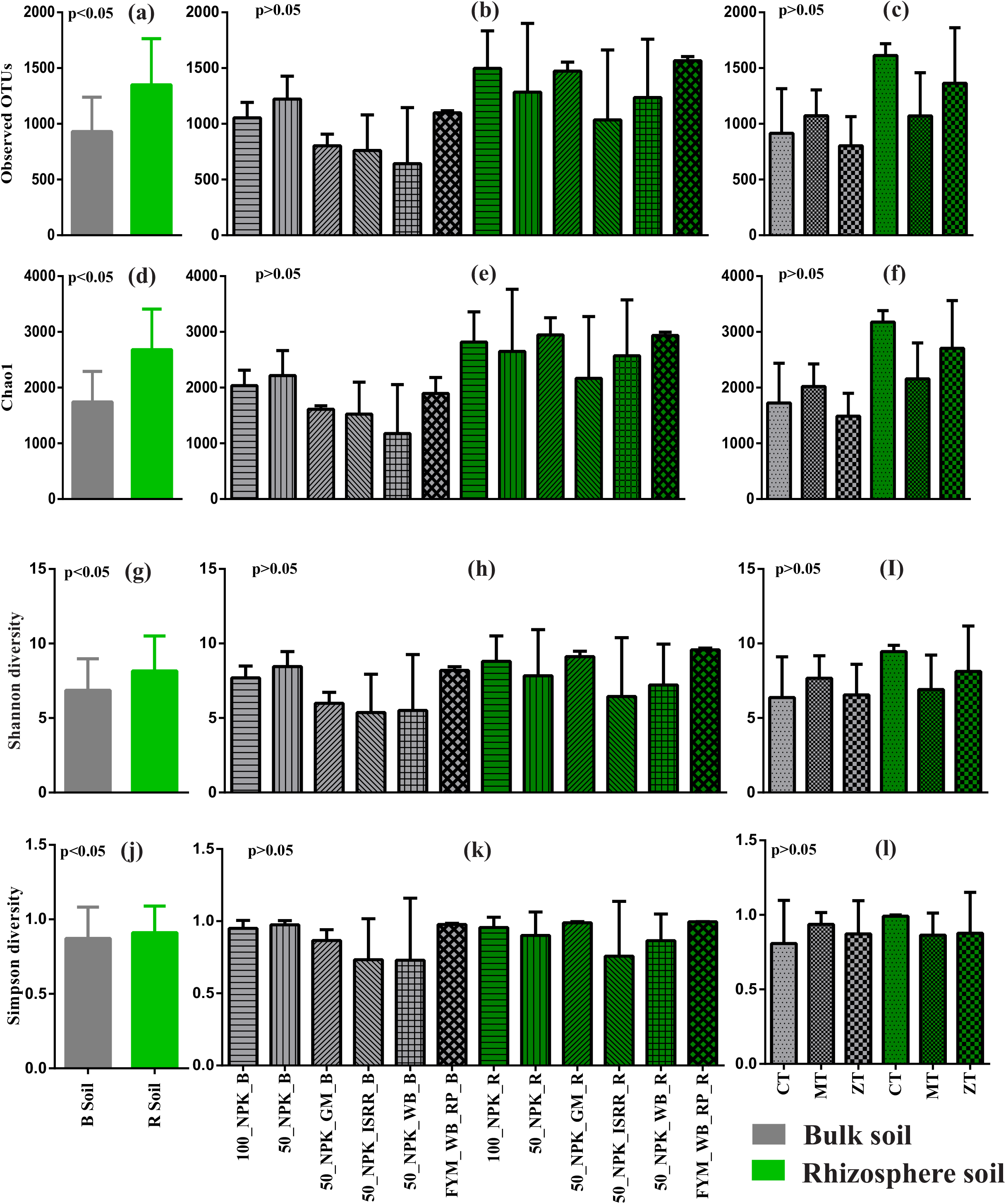
Bar charts with standard error bars illustrating the values of alpha diversity measures, Observed OTUs (a-c) Chao1 (d-f), Shannon (g-i) and Simpson (j-l) in the bulk soil and pea rhizosphere samples from different tillage and residue management practices.

The beta diversity analysis based unweighted unifrac PCoA showed clustering of the bulk soil and rhizosphere soil sample into two distinct clusters (**Fig. 2**). The total variation explained by first was PCoA 34.8% (22% for PCoA1 and 12.8% for PCoA2), wherein all the bulk soil samples were on the positive side of the PC2 except B15 and B03 while all the rhizosphere soil samples were on the negative side of the axis (**Fig. 1e**).

**Fig. 2.**
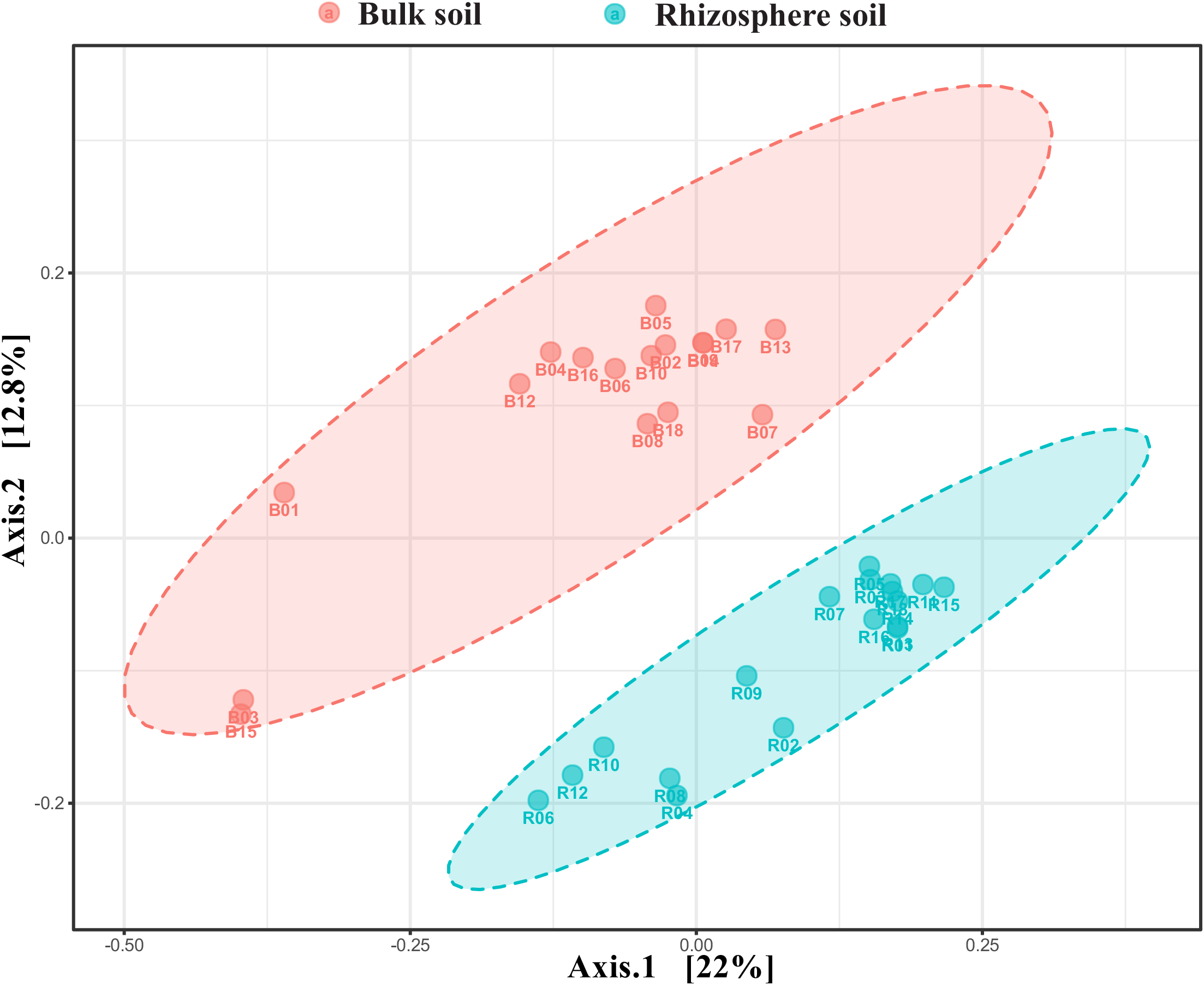
PCoA bi-plot based on relative abundance of bacterial OTUs exhibiting the beta diversity among the bulk soil and pea rhizosphere samples.

### 3.2. Taxonomic composition of bacterial communities

Total 71 bacterial phyla were detected in the bulk soil and rhizosphere samples in which, Proteobacteria (32.5%), Firmicutes (29.4%), Acidobacteria (9.03%), Actinobacteria (7.2%), Chloroflexi (4.7%), Nitrospirae (3.1%), Verrucomicrobia (2.7%), Thaumarchaeota (2.6%), Bacteroidetes (2.1%) and Planctomycetes (1.03%) were highly abundant and constituted 95% of the overall bacterial community (**Fig. 3a**). Group’s specific abundance pattern was seen for Proteobacteria and Firmicutes wherein, Firmicutes were highly abundant in bulk soil (~41.7%) than rhizosphere (~17.8%) while, Proteobacteria were highly abundant in rhizosphere (~43.9%) in comparison to bulk soil (~18.6%) samples (**Fig. 3a**).

**Fig. 3.**
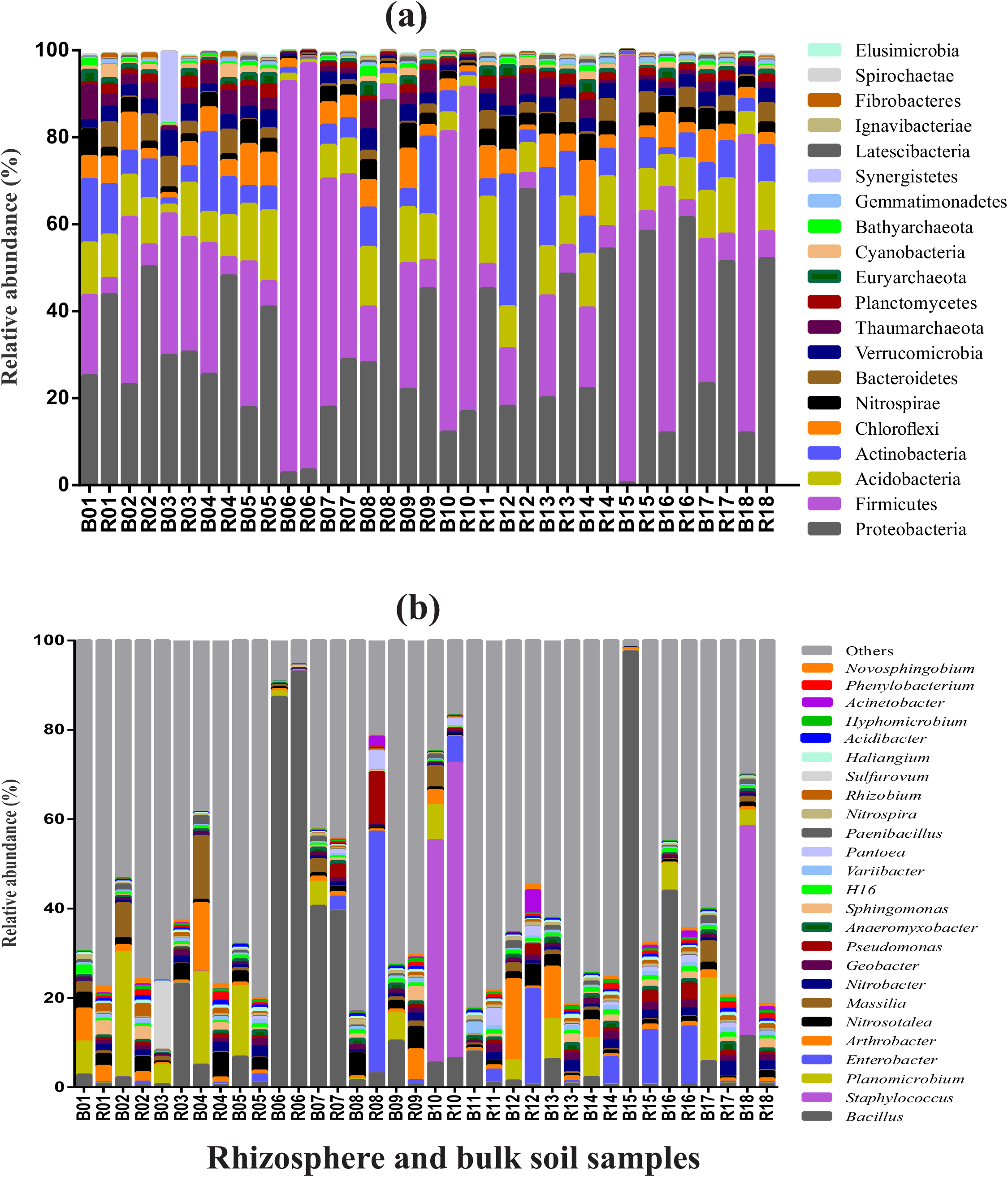
Relative abundance of bacteria in the bulk soil and pea rhizosphere samples at the phylum (a) and genus (b) levels.

Bacterial genera *Bacillus, Staphylococcus, Planomicrobium, Enterobacter, Arthrobacter, Nitrosotalea, Massilia, Nitrobacter, Geobacter, Pseudomonas, Anaeromyxobacter, Sphingomonas*, candidates genus *H16, Variibacter, Pantoea, Paenibacillus, Nitrospira, Rhizobium, Sulfurovum, Haliangium, Acidibacter, Phenylobacterium, Hyphomicrobium, Acinetobacter* and *Novosphingobium* were found dominant with a collective abundance more than 0.5% (**Fig. 3b**). A high abundance of *Planomicrobium* and *Arthrobacter* was found in the bulk soil samples and *Enterobacter* in the rhizosphere soil samples. *Arthrobacter* was highly abundant (in a range of 7% to 18%) in the samples B01, B04, B12, B13 and R09 (6.9%), while remaining samples showed a lesser abundance (0.1 to 4%). Notably, for bulk soil sample B06, B07, B15, B16, B18, and rhizosphere samples R03, R06 and R07 have a high abundance of *Bacillus* (**Fig. 3b**).

Comparative analysis at higher (phylum) and lower (genus) taxonomic ranks revealed the presence of differentially abundant bacterial taxa in bulk soil and rhizosphere soil samples. Phyla Proteobacteria and Bacteroidetes were abundant in rhizosphere soil while Firmicutes, Chloroflexi, Nitrospirae, and Planctomycetes were highly abundant in bulk soil (t-test, p<0.05, Benjamini-Hochberg FDR corrections) (**Fig. 4a**). The abundance of genera *Rhizobium, Pseudomonas, Pantoea, Paenibacillus, Nitrobacter, Enterobacter*, and *Sphingomonas* were significantly higher in rhizosphere soils, while *Massilia, Paenibacillus* and *Planomicrobium* were high in bulk soils (t-test, p<0.05) (**Fig. 4b**). Overall, the presence of 917 bacterial genera was observed in both rhizosphere and bulk soil samples. Among 917 genera 60% (551 nos.) were shared between the bulk soil and rhizosphere soil samples, while ~29% (267 nos.) were unique to the rhizosphere, and ~11% (99 nos.) genera were exclusive to bulk soil sample (**Fig. 4c**).

**Fig. 4.**
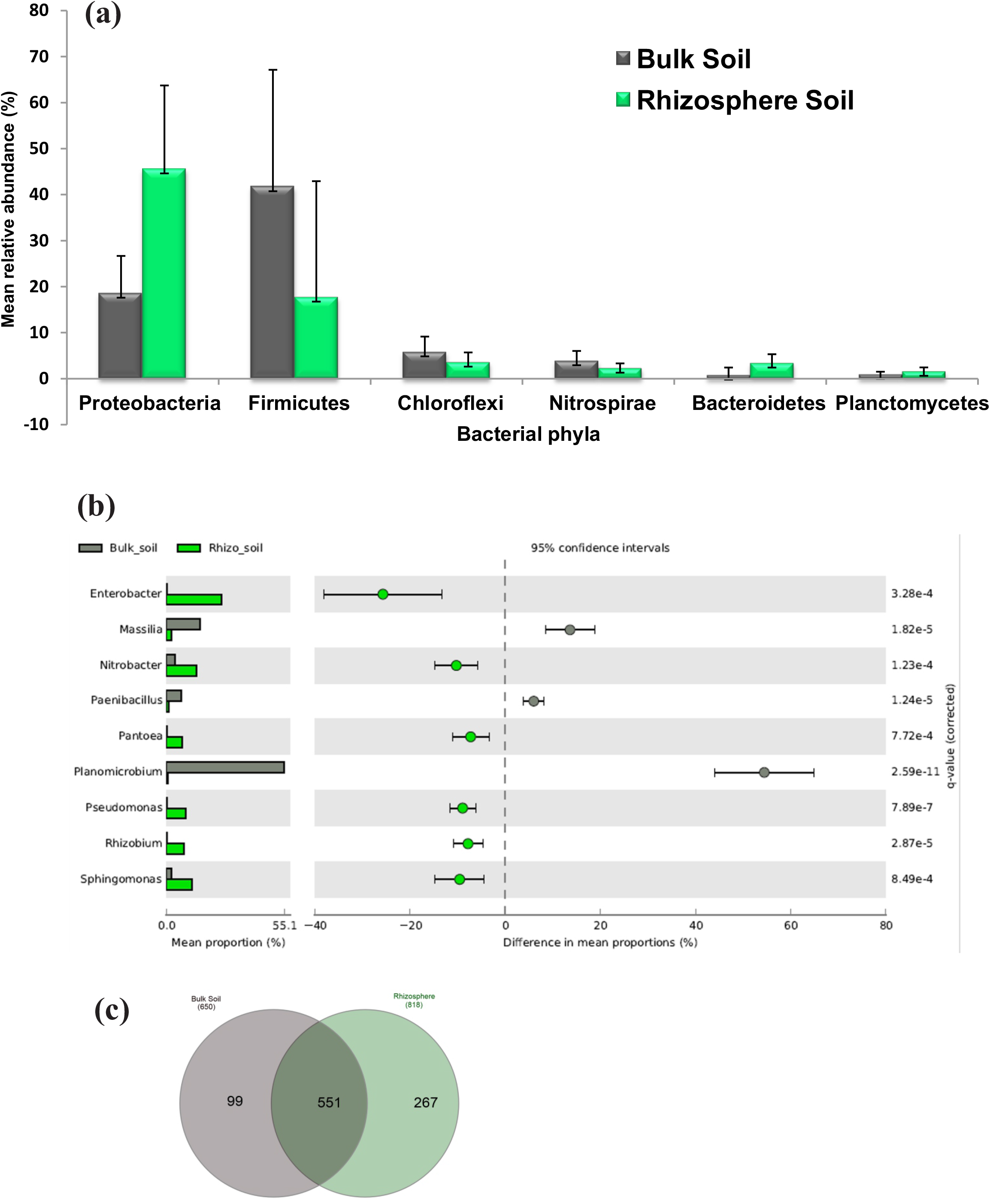
Differentially abundant bacterial taxa across the bulk soil and pea rhizosphere samples. A) At the phylum level. B) At the genera level C) Venn diagram representing the number of shared and unique OTUs in the bulk soil and rhizosphere soil samples.

### 3.3. Quantitative analysis of bacterial community

The average total bacterial abundance in the rhizosphere (8.9 x 10^9^ copies/gm) and bulk soil (9.42 x 10^9^ copies/gm) samples was statistically not different from each other (**Fig. 5a**). Quantitative enumeration of the genus Firmicutes depicted the significantly higher copy number (3.76 x 10^9^ copies/gm) in the bulk soil samples in comparison to the rhizosphere samples (1.78 x 10^9^ copies/gm) (**Fig. 5b**). The mean relative abundance of Firmicutes detected in 16S rRNA gene amplicon sequencing data was ~18% in the rhizosphere samples and ~42% in the bulk soil samples (**Fig. 5c**).

**Fig. 5.**
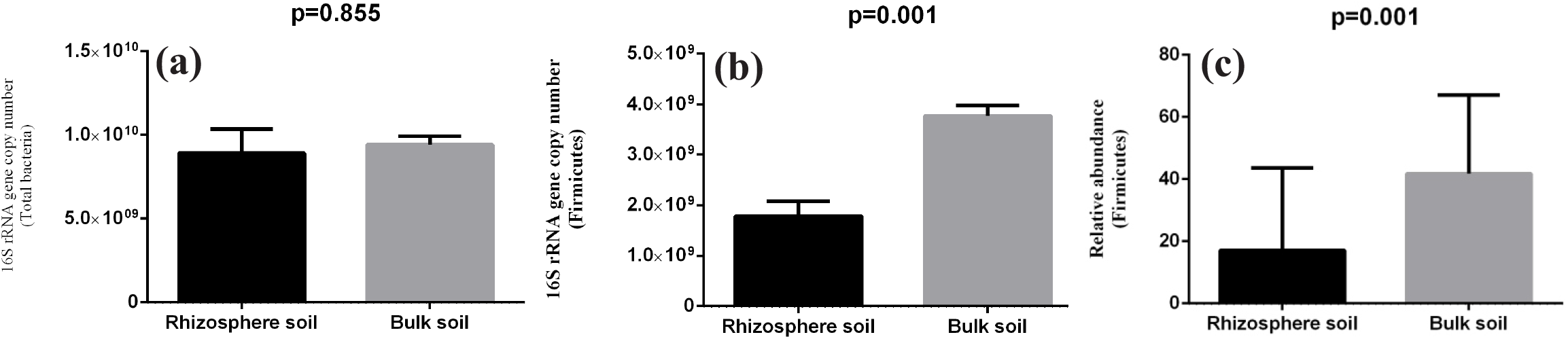
Abundance of total bacteria and phylum Firmicutes in the bulk soil and rhizosphere soil samples. A) Total bacterial abundance in the bulk soil and rhizosphere soil samples, B) qPCR based and C) NGS based relative abundance of phylum Firmicutes in the bulk soil and rhizosphere soil samples.

### 3.4. Effect of soil management practices on the bulk soil and pea rhizosphere microbiome

Bulk soil microbial community showed the differential abundance of *Pseudolabrys, Roseiarcus*, and *Tumebacillus* across the residue management practices (ANOVA, p<0.05). The abundance of *Pseudolabrys* and *Tumebacillus* was high in samples from 100% NPK and organic treatments, while *Roseiarcus* was abundant in 50% NPK and organic treatment field samples (**Fig. 6a**). Similarly, in the bacterial community analysis of the rhizosphere soil bacterial genera *Anaerolinea, Hydrogenispora*, and *Syntrophorhabdus* were differentially abundant across the groups and having higher abundance in the organic treatment farm (**Fig. 6b**).

**Fig. 6.**
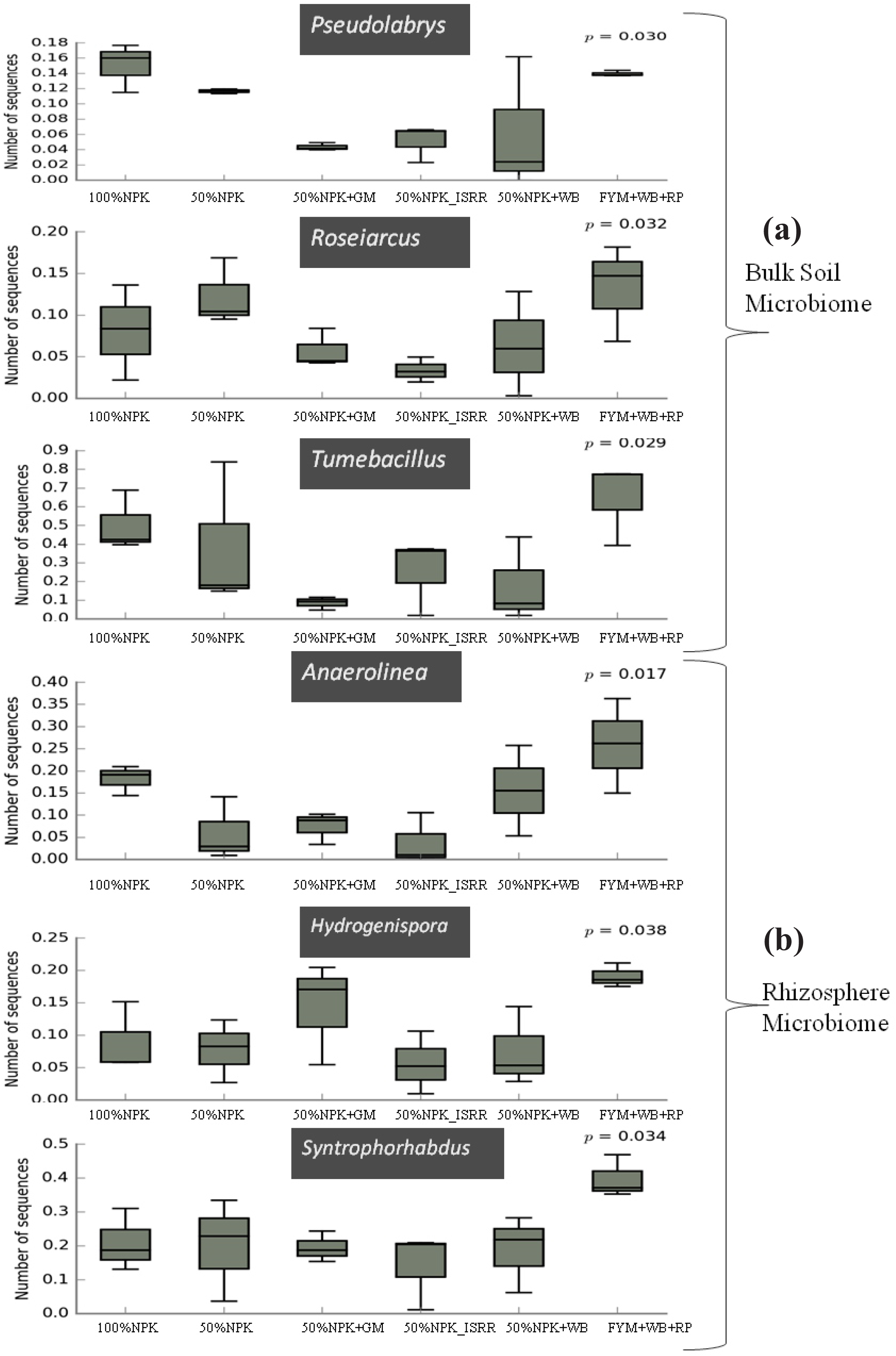
Differentially abundant bacterial taxa detected across the different residue management practice groups in the a) bulk soil and b) pea rhizosphere samples.

Three different soil management practices revealed enrichment of specific OTUs in the bulk soil and rhizospheric soil (ANOVA, p<0.05). Among the bulk soil samples 12 OTUs belonging to four taxa were differentially abundant in conventional tillage, 66 OTUs from 32 taxa in minimum tillage and only six OTUs from four taxa in zero tillage (**Fig. 7a**). Similarly, in the rhizosphere soil samples 295 OTUs of 96 taxa, 17 OTUs of the 13 taxa and 82 OTUs of the 37 taxa were significantly enriched in the conventional tillage, minimum tillage and zero tillage, respectively (**Fig. 7b**). At the genera-level bacterial community analysis across three tillage treatment fields, 11 bacterial genera showed differential abundance (**Table 2**). All these genera showed higher abundance in the conventional tillage fields.

**Fig. 7.**
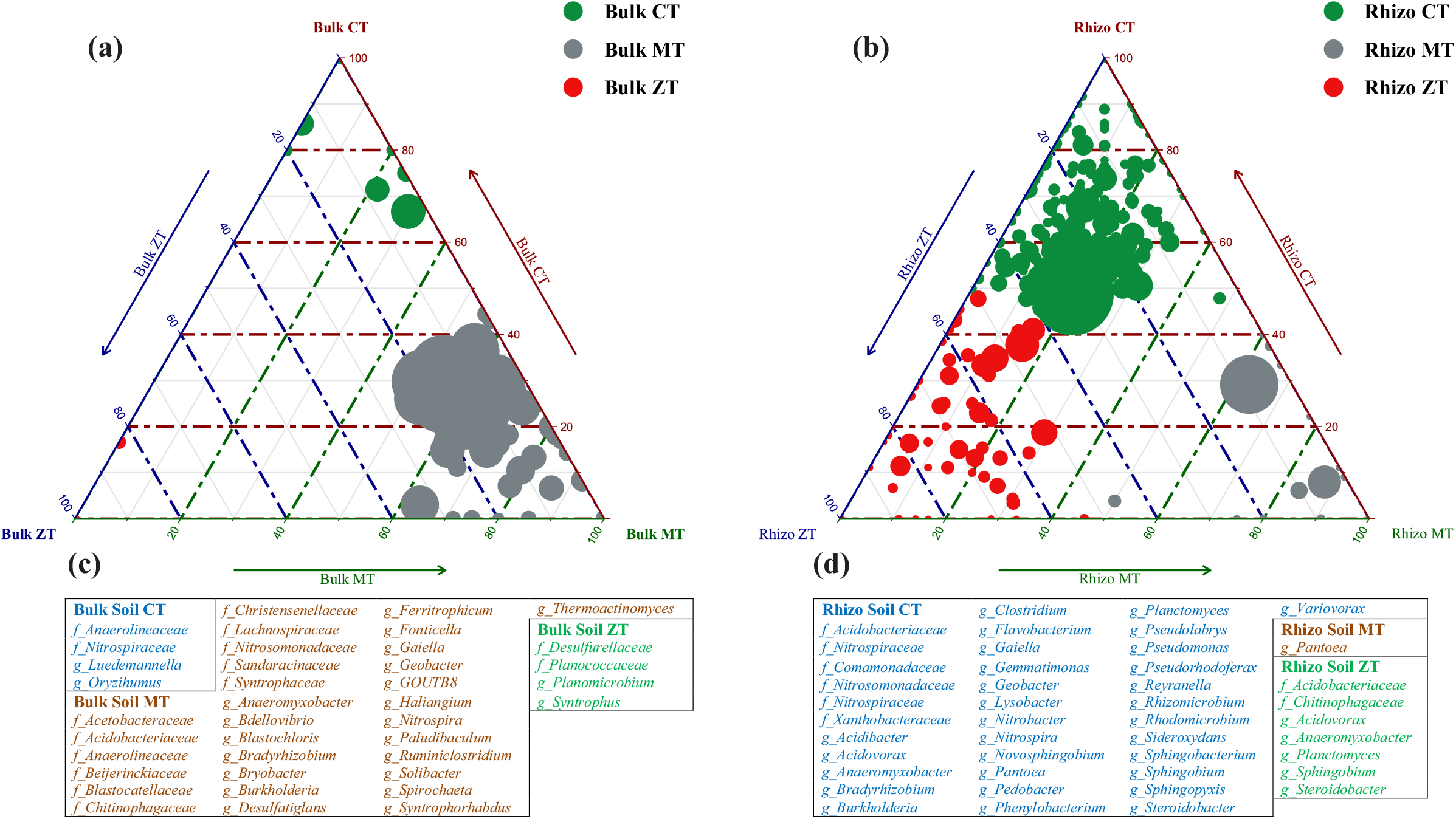
Ternary plot depicting the number of OTUs enriched in the zero tillage, minimum tillage and conventional tillage plots in the a) bulk soil and b) pea rhizosphere samples. Each circle depicts one individual OTU. The size of the circle reflects its relative abundance. Family or genus level taxonomy of the OTUs specifically enriched in the bulk soil (c) and rhizosphere soil (d) in respective tillage treatments. For the rhizosphere soil samples taxonomy of OTUs having abundance more than 0.05% is represented.

**Table 2:** Differentially abundant bacterial genera across the three different tillage treatments practices

| **Genera** | **Zero tillage**<br>**(Mean ± STD)** | **Conventional tillage**<br>**(Mean ± STD)** | **Minimum tillage**<br>**(Mean ± STD)** | **p-values**<br>**(ANOVA)** |
| --- | --- | --- | --- | --- |
| *Flavobacterium* | 0.989 ± 0.684 | 2.197 ± 0.428 | 0.523 ± 0.54 | 0.001 |
| *Lysobacter* | 0.639 ± 0.426 | 1.194 ± 0.354 | 0.269 ± 0.174 | 0.002 |
| *Pseudolabrys* | 1.071 ± 0.533 | 1.298 ± 0.158 | 0.345 ± 0.36 | 0.003 |
| *Rhodomicrobium* | 0.437 ± 0.253 | 0.888 ± 0.18 | 0.299 ± 0.264 | 0.003 |
| *Pedomicrobium* | 0.297 ± 0.164 | 0.578 ± 0.197 | 0.194 ± 0.15 | 0.008 |
| *Acidothermus* | 0.421 ± 0.233 | 0.836 ± 0.418 | 0.2 ± 0.127 | 0.01 |
| *Bryobacter* | 1.285 ± 0.663 | 1.54 ± 0.32 | 0.473 ± 0.463 | 0.011 |
| *Anaeromyxobacter* | 1.763 ± 1.184 | 3.445 ± 1.398 | 1.107 ± 0.885 | 0.018 |
| *Rhizomicrobium* | 0.72 ± 0.645 | 1.179 ± 0.423 | 0.274 ± 0.224 | 0.025 |
| *Nitrobacter* | 4.096 ± 2.36 | 5.912 ± 1.874 | 2.104 ± 1.812 | 0.031 |
| *Sideroxydans* | 0.401 ± 0.366 | 0.629 ± 0.121 | 0.195 ± 0.196 | 0.047 |

### 3.5. Predictive functional analysis of the microbial community

The functional contributions of the bacteria were predicted based on OTUs, and the results revealed the presence of 398 different functional classes. Among these classes, transporters, two-component system, secretion system, ABC transporters, transcription factors, peptidases, ribosome, methane metabolism, quorum sensing, and bacterial motility proteins were the top 10 highly represented classes. The presence of 6596 KEGG orthologues (KO) was predicted across all samples, belonging to metabolism, environmental information processing, cellular processes, human diseases, genetic information processing, and organismal systems. Principle component analysis based on the highly abundant functions clustered the bulk soil and rhizosphere samples into separate groups (**Fig 8**). Majority of the rhizosphere samples are placed on the negative side of PC1 and positive side of PC2, while most of the bulk samples were on the positive side of PC1 and negative side of PC2 (**Fig 8**). Bacterial genes associated with iron complex outer membrane receptor protein, cobalt-zinc-cadmium resistance protein (CzcA), RNA polymerase sigma-70 factor, ribonuclease E, translation initiation factor IF-2, serine/threonine protein kinase, hydrophobic/amphiphilic exporter-1, beta-glucosidase and multiple sugar transport system permease protein showed high negative values for PC1 and positive for PC2 (Table S3). Rhizosphere samples revealed high abundance of genes involved in nitrogen fixation (*NifQ*) by 64%, enterobactin (siderophore) production by 39%, plant hormone IAA production (tryptophan 2-monooxygenase) by 10%, phosphate metabolism (inorganic phosphate transporter) by 40% in comparison to bulk soil samples (**Supp file S1**).

**Fig. 8.**
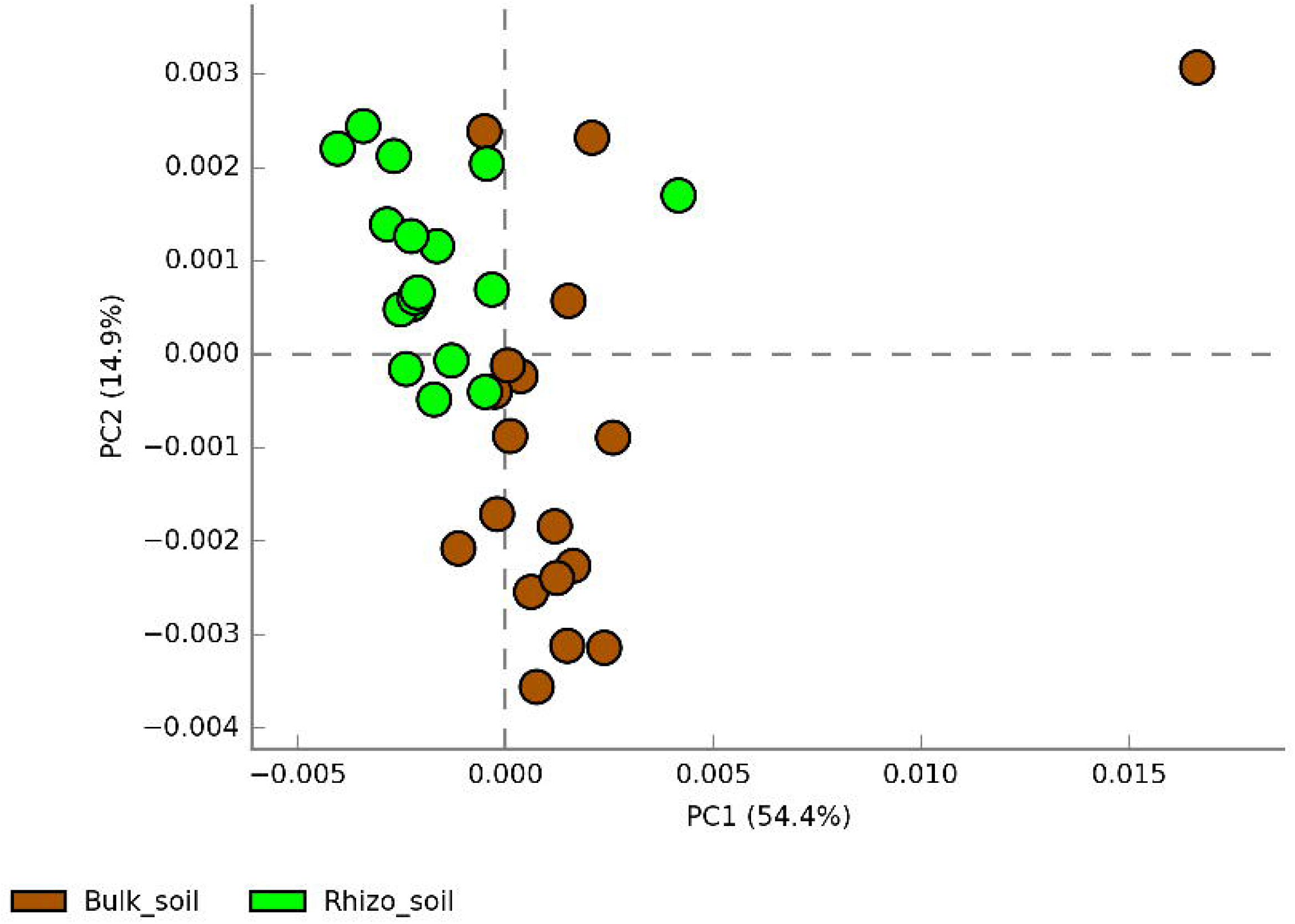
PCA bi-plot based on relative abundance predictive metabolic potentials of bacterial communities in the bulk soil and pea rhizosphere samples.

## 4. Discussion

Agriculture is a continually improving system to make it resilient and sustainable for the long term and meet the ever-increasing human requirements. Conservation agriculture offers a framework to improve soil structure, save water, enhance soil nutrient supply and cycling, increase yield, and maintain soil biodiversity. This study was designed to investigate the effect of long-term exposure to various tillage and residue management practices on the bacterial community structures of the bulk soils, and how pea plant shapes the rhizosphere communities. Soil bacterial diversity is known to be influenced by the numerous environmental factors including soil pH (Lauber et al., 2009; Cho et al., 2016; Sánchez-Cañizares et al., 2017), chemistry of soil organic matter (Lupatini et al., 2017), plant species grown in the soil (Yamamoto et al., 2018), secretion of different root exudates by the plant species (Sasse et al., 2017; Huang et al., 2019) and processing of the soil before cultivation of the plants (Moreno-Espíndola et al., 2018). Majority of the soils in North-East regions of India are acidic (pH = 5.0 to 6.0) and rich in organic carbon, which makes them deficient in available phosphorous, medium to low in available potassium, and highly rich in iron and aluminum (Thakuria et al., 2016). Our results showed the dominance of Proteobacteria, Firmicutes, Acidobacteria and Actinobacteria in both bulk and rhizosphere soils (Fig 3), and are in agreement to the previous report on the dominance of copiotrophic microorganisms like Proteobacteria, Firmicutes, and Actinobacteria in rich organic environments, and soils with low pH harbour more Acidobacteria (Fierer et al., 2007).

Significant differences were observed in the overall alpha diversity in the rhizosphere and bulk soil samples (**Fig. 1**), demonstrating higher species abundance and evenness (based on Shannon and Simpson indices) in rhizosphere samples. There was no significant difference in bacterial richness and evenness among the different tillage and residue management treatments (P□>□0.05) in both rhizosphere and bulk soils (**Fig. 1**), indicating that plant rhizosphere effect is the key driver for alpha diversity. Plants can alter the microbial communities by secreting a variety of nutrients and bioactive molecules into the rhizosphere (Hu et al., 2018; Huang et al., 2019). The enrichment of specific OTUs in the pea rhizosphere leading to the increased diversity was further confirmed, as all the rhizosphere samples were grouped in a small cluster, in comparison to a loose clustering of bulk soil samples in the PCoA biplot (**Fig. 2**). The built-up of homogeneous bacterial communities in most of the rhizosphere samples can be attributed to the selection pressure of the pea roots, which continually release a large number of border cells and mucilage (Ropitaux et al., 2019), and pose a strong rhizosphere effect on the bacteria. Impact of crop plant on the diversity of rhizosphere microbes has also been observed for different crops including barley, cotton, maize and wheat (Yurgel et al., 2018; Babin et al., 2019; Kerdraon et al., 2019).

Majority of members of microbial communities in the host plant are horizontally acquired from the surrounding environment, and the soil is the main reservoir of a plant rhizosphere microbiome (Wagner et al., 2016; Sánchez-Cañizares et al., 2017). Our results on pea rhizosphere and bulk soils are consistent with this, as 551 (60%) genera of 917 were common in bulk and rhizosphere soil samples (**Fig. 4c**). Dominance of Proteobacteria recorded in pea rhizosphere samples, with significant increase in the abundance of genera including *Pseudomonas, Rhizobium, Pantoae, Enterobacter* and *Sphingomonas* known for plant growth promoting attributes, was in agreement with the previous studies on the rhizosphere microbiome analysis (Mendes et al., 2013; Weinert et al., 2011; Yurgelet al., 2018; Goss-Souza et al., 2019). Pea-Rhizobium symbiosis is well documented in Indian soils (Rahi et al., 2012), hence increase in the abundance of *Rhizobium* in pea rhizosphere was expected. Higher abundance of *Pseudomonas* and *Sphingomonas* was also reported in the rhizospheres of crop plants like lettuce, wheat and maize (Schreiter et al., 2014; Kerdraon et al., 2019). In addition to Proteobacteria, a significantly higher abundance of Bacteroidetes and Planctomycetes was also recorded in the rhizosphere soil in comparison to bulk soil samples (**Figs. 3-4**), positive correlations with increased abundance of Bacteroidetes and Planctomycetes have been observed to increase in organic carbon and phosphorus concentrations, respectively in rhizosphere of soybean (Goss-Souza et al., 2019). The genus *Nitrobacter* was at higher abundance in pea rhizosphere samples than bulk soils (Fig 4b), suggesting its enrichment by host plant as *Nitrobacter* converts nitrite to nitrate making nitrogen more readily available to the host plant (Richardson et al., 2009; Hubbard et al., 2018).

The abundance of Firmicutes represented by members of genera like *Bacillus*, *Staphylococcus* and, *Planomicrobium*, witness a significant decrease in pea rhizosphere in comparison to bulk soil samples (**Figs. 3-5**), previous studies have also indicated negative correlation of plant growth and high abundance of Firmicutes in the soil (Zhang et al., 2014; Kumar et al., 2018). The consistently higher relative proportion of genus *Planomicrobium* in the majority of bulk soil samples (**Fig. 3**) also confirms its dominance in soils (Hue et al., 2019). The abundance of bacterial phyla Chloroflexi and Nitrospirae was significantly higher in bulk soil to rhizosphere samples (**Fig. 4a**), which is in agreement with the previous study on the impact of land use intensity and plant functional identity on microbial communities (Schöps et al., 2018). The members of Chloroflexi are known for the decomposition of organic matter (Wu et al., 2016), which is expected to be high in the bulk soils because of different organic inputs and paddy crop residue in most of the treatments. Nitrospirae are slow-growing microbial K-strategists adapted to low substrate concentrations (Daims et al., 2015), and could not cope up other fast-growing bacterial communities in the nutrient-rich pea rhizosphere environment. Nitrospirae are involved in nitrification process (oxidation of nitrite to nitrate) and have been reported to be dominant in N-fertilized treatment in paddy soil (Kumar et al., 2018). Significantly high abundance of *Massilia* and *Paenibacillus* was observed in bulk soil than rhizosphere samples (**Fig. 4b**), which can be attributed to presence of high cellulosic biomass in most of the treatment leading to the selection of the members of genera with potential to degrade cellulose (Ofek et al., 2012; Grady et al., 2016).

Minor differences were observed in the bacterial community composition in response to residue management treatments in both bulk and rhizosphere soils (**Fig. 6**), exhibiting the complex responses of microbial communities to fertilizer applications (Hartmann et al., 2015). Differential abundance of members of genera *Pseudolabrys, Roseiarcus* and *Tumebacillus* were recorded in bulk soils samples, and *Anaerolineae, Hydrogenispora* and *Syntrophorhabdus* in rhizosphere samples across the residue management treatments (**Fig. 6**). The higher proportion of *Pseudolabrys* in 100% NPK and organic treatments, indicate its specific enrichment in these treatments, *Pseudolabrys* is a member of Rhizobiales, reported from soil and lettuce rhizosphere (Cipriano et al., 2016). *Roseiarcus* is a methanotroph isolated for acidic peat soil (Kulichevskaya et al., 2014), the function of this genera is not well understood and was abundant in the organic treatment (**Fig. 6**). Increased abundance of *Tumebacillus* in organic treatment in the acidic soils of Meghalaya, suggesting that in acidic soils aluminum stress activates *Tumebacillus* to improve aluminum stress tolerance in host plants (Lian et al., 2019). Among the rhizosphere sample significant increase has been observed in the relative abundance of *Anaerolineae*, *Hydrogenispora* and *Syntrophorhabdus* in the organic treatment (**Fig. 6**), the organic treatment increase soil carbon content and nutrients, which are the key drivers for soil microbial community (Fierer et al., 2007; Bei et al., 2018). All of these genera are anaerobic bacteria capable of decomposing diverse carbon sources under anoxic environments and have identified as the unique core taxa for rice soils (Jiao et al., 2019).

Though pea was cultivated under zero-tillage, influence of previous tillage treatments during rice cultivation was observed on the prevalence of few specific OTUs (**Fig. 7**), suggesting the plausible effect of tillage treatments on the bacterial community structure. This can be attributed to the fact that tillage practices alter soil bulk density, pore structure, water availability, soil organic carbon, etc (Zhang et al., 2018). However, the impact to tillage treatments was not pronounced, and only 0.56% OTUs in the bulk soil, and 2.60% OTUs in the rhizosphere soil were enriched across various tillage treatments (**Fig. 7**). Increased abundance of OTUs assigned to *Acidothermus, Anaeromyxobacter, Bryobacter, Flavobacterium, Lysobacter, Nitrobacter, Pedomicrobium, Rhizomicrobium, Pseudolabrys, Rhodomicrobium*, and *Sideroxydans* was recorded in the rhizosphere soil samples in the conventional tillage in comparison to minimal and zero tillage (**Table 2**). Severe soil disturbance in conventional tillage accelerates soil organic matter oxidation hence expected to enrich a few members of bacteria (Ding et al., 2011).

Higher abundance of genes related to nitrogen fixation, phytohormone and siderophore production, phosphate metabolism, etc. in the rhizosphere soil (**Supp file S2**) substantiate our earlier conclusions on the selection of bacterial communities with plant growth promoting potential in the rhizosphere (**Fig. 3**). Principle component analysis based on the highly abundant functions clustered the bulk soil and rhizosphere samples into separate groups, indicating selection of specific bacterial communities in rhizosphere based on their putative functions (**Fig. 8**). In addition to high abundance of plant growth promotion genes, bacterial genes associated with iron complex outer membrane receptor protein, cobalt-zinc-cadmium resistance protein (CzcA), RNA polymerase sigma-70 factor and ribonuclease E were also abundant in rhizosphere soil as compared to the bulk soil (**Supp file S2**), revealing the possible role of bacterial communities in abiotic stress (low pH and high aluminum and iron toxicity) amelioration.

## 6. Conclusions

Our work showed that pea plant is the most dominating selection factor shaping the microbial communities under diverse residue management and tillage treatments. Soil management and residue management practices also affect the bacterial community structure in both bulk soil and pea rhizosphere. However, their influence was not significantly different. Enrichment of bacterial taxa known for plant growth promotion attributes and removal of toxic elements from soil was recorded in the rhizosphere indicating selection of rhizosphere communities by the plant to meet its requirements of nutrient uptake and combating stress. Predictive functional analysis also revealed the plausible enrichment of plant growth promoting and stress tolerance genes in the pea rhizosphere.

## Acknowledgments

Authors acknowledge the Director, NCCS, Pune, and financial support by the SERB-DST (YSS/2015/000149), and Department of Biotechnology (BT/IN/Indo-US/Foldscope/39/2015).

## References

Aßhauer, K. P., Wemheuer, B., Daniel, R., and Meinicke, P. (2015). Tax4Fun: predicting functional profiles from metagenomic 16S rRNA data. Bioinformatics 31, 2882–2884. doi: 10.1093/bioinformatics/btv287

Babin, D., Deubel, A., Jacquiod, S., Sørensen, S. J., Geistlinger, J., Grosch, R., et al. (2018). Impact of long-term agricultural management practices on soil prokaryotic communities. Soil Biol. Biochem. 129, 17–28. doi: 10.1016/j.soilbio.2018.11.002

Bei, Y. Zhang, T. Li, P. Christie, X. Li, J. Zhang. (2018). Response of the soil microbial community to different fertilizer inputs in a wheat-maize rotation on a calcareous soil Agric. Ecosyst. Environ. 260, 58–69.

Cho, S.-J., Kim, M.-H., and Lee, Y.-O. (2016). Effect of pH on soil bacterial diversity. J. Ecol. Environ. 40, 10. doi: 10.1186/s41610-016-0004-1

Cipriano, M. A., Lupatini, M., Lopes-Santos, L., Da Silva, M. J., Roesch, L. F., Destéfano, S. A., et al. (2016). Lettuce and rhizosphere microbiome responses to growth promoting Pseudomonas species under field conditions. FEMS Microbiol. Ecol. 92, fiw197. doi: 10.1093/femsec/fiw197

Daims, H., Lebedeva, E. V., Pjevac, P., Han, P., Herbold, C., Albertsen, M., et al., (2015). Complete nitrification by Nitrospira bacteria. Nat. 528, 504–509. doi: 10.1038/nature16461

Degrune, F., Theodorakopoulos, N., Colinet, G., Hiel, M-P., Bodson, B., Taminiau, B., Daube, G., Vandenbol, M., Hartmann, M. (2017). Temporal Dynamics of Soil Microbial Communities below the Seedbed under Two Contrasting Tillage Regimes. Front. Microbiol. 8, 1127. doi: 10.3389/fmicb.2017.01127

Dhariwal, A., Chong, J., Habib, S., King, I. L., Agellon, L. B., and Xia, J., (2017). Microbiome analyst: a web-based tool for comprehensive statistical, visual and meta-analysis of microbiome data. Nucleic Acids Res. 45, W180–W188. doi: 10.1093/nar/gkx295

Ding, X., Zhang, B., Zhang, X., et al., (2011). Effects of tillage and crop rotation on soil microbial residues in a rainfed agroecosystem of northeast China Soil. Tillage Res., 114 (2011), pp. 43–49

Edgar, R. C. (2010). Search and clustering orders of magnitude faster than BLAST. Bioinformatics 26, 2460–2461. doi: 10.1093/bioinformatics/btq461

Fadrosh, D. W., Ma, B., Gajer, P., Sengamalay, N., Ott, S., Brotman, R. M., et al. (2014). An improved dual-indexing approach for multiplexed 16S rRNA gene sequencing on the Illumina MiSeq platform. Microbiome 2, 6. doi: 10.1186/2049-2618-2-6

Fierer, N., Bradford, M. A., and Jackson, R. B., (2007). Toward an ecological classification of soil bacteria. Ecol. 88, 1354–1364. doi: 10.1890/05-1839

Gan, Yantai et al. (2015). Diversifying crop rotations with pulses enhances system productivity. Sci. Rep. 5, 14625. doi: 10.1038/srep14625

Godfray, H. C. J., and Garnett, T. (2014). Food security and sustainable intensification. Philos. Trans. R. Soc. Lond. B Biol. Sci. 369(1639), 20120273. doi: 10.1098/rstb.2012.0273.

Goss-Souza, D., Mendes, L.W., Borges, C.D., Rodrigues, J.L.M., Tsai, S.M. (2019) Amazon forest-to-agriculture conversion alters rhizosphere microbiome composition while functions are kept. FEMS Microbiol Ecol. 95(3), fiz009, doi: 10.1093/femsec/fiz009.

Grady, E. N., MacDonald, J., Liu, L., Richman, A., and Yuan, Z. C. (2016). Current knowledge and perspectives of Paenibacillus: a review. Microb. Cell Fact. 15, 203. doi: 10.1186/s12934-016-0603-7

Guo, X., Xia, X., Tang, R., Zhou, J., Zhao, H., and Wang, K. (2008). Development of a real-time PCR method for Firmicutes and Bacteroidetes in faeces and its application to quantify intestinal population of obese and lean pigs. Lett. Appl. Microbiol. 47, 367–373. doi: 10.1111/j.1472-765X.2008.02408.x

Hartmann, M., Frey, B., Mayer, J., Mäder, P., Widmer, F. (2014). Distinct soil microbial diversity under long-term organic and conventional farming. ISME J. 9(5), 1177–94. doi: 10.1038/ismej.2014.210.

Herridge, D. F., Peoples, M. B., Boddey, R. M. (2008). Global inputs of biological nitrogen fixation in agricultural systems. Plant Soil. 311, 1–18. doi: 10.1007/s11104-008-9668-3.

Hogenhout, S. A., and Loria, R. (2008). Virulence mechanisms of Gram-positive plant pathogenic bacteria. Curr. Opin. Plant Biol. 11, 449–456. doi: 10.1016/j.pbi.2008.05.007

Hu, L., Robert, C. A. M., Cadot, S., Zhang, X., Ye, M., Li, B., et al. (2018). Root exudate metabolites drive plant-soil feedbacks on growth and defense by shaping the rhizosphere microbiota. Nat. Commun. 9, 2738. doi: 10.1038/s41467-018-05122-7

Huang, A.C., et al. (2019) A specialized metabolic network selectively modulates *Arabidopsis* root microbiota. Sci. 364(6440), eaau6389. DOI: 10.1126/science.aau6389

Hubbard, C. J., Li, B., McMinn, R., et al. (2019). The effect of rhizosphere microbes outweighs host plant genetics in reducing insect herbivory. Mol Ecol. 28(7), 1801–1811. doi: 10.1111/mec.14989.

Hui N, Grönroos M, Roslund MI, Parajuli A, Vari HK, Soininen L, Laitinen OH, Sinkkonen A and The ADELE Research Group (2019). Diverse Environmental Microbiota as a Tool to Augment Biodiversity in Urban Landscaping Materials. Front. Microbiol. 10, 536. doi: 10.3389/fmicb.2019.00536

Kerdraon, L., Balesdent, M.H., Barret, M., Laval., Suffert, F. (2019). Crop Residues in Wheat-Oilseed Rape Rotation System: a Pivotal, Shifting Platform for Microbial Meetings. Microb. Ecol. 77(4), 931–945. doi: 10.1007/s00248-019-01340-8.

Kerdraon, L., Balesdent, MH., Barret, M. et al. (2019) Crop Residues in Wheat-Oilseed Rape Rotation System: a Pivotal, Shifting Platform for Microbial Meetings. Microb. Ecol. 77, 931. https://doi.org/10.1007/s00248-019-01340-83

Kulichevskaya, I. S., Danilova, O. V., Tereshina, V. M., Kevbrin, V. V., and Dedysh, S. N., (2014). Descriptions of *Roseiarcus fermentans* gen. nov., sp. nov., a bacteriochlorophyll a-containing fermentative bacterium related phylogenetically to alphaproteobacterial methanotrophs, and of the family Roseiarcaceae fam. nov. Int. J. Syst. Evol. Microbiol. 64, 2558–2565. doi: 10.1099/ijs.0.064576-0

Kumar, U., Nayak, A.K., Shahid, M. et al. (2018) Continuous application of inorganic and organic fertilizers over 47 years in paddy soil alters the bacterial community structure and its influence on rice production. Agric. Ecosyst. Environ. 262, 65–75. https://doi.org/10.1016/j.agee.2018.04.016

Laik R., Sharma S., Idris M., Singh A.K., Singh S.S., Bhatt B.P., Saharawat Y.S., Humphreys E., Ladha J.K. (2014). Integration of conservation agriculture with best management practices for improving system performance of the rice-wheat rotation in the Eastern Indo-Gangetic Plains of India. Agric. Ecosyst. Environ. 195, 68–82. https://doi.org/10.1016/j.agee.2014.06.001

Lauber, C. L., Hamady, M., Knight, R., and Fierer, N. (2009). Pyrosequencing-based assessment of soil pH as a predictor of soil bacterial community structure at the continental scale. Appl. Environ. Microbiol. 75, 5111–5120. doi: 10.1128/AEM.00335-09

Lian, T., Ma, Q., Shi, Q. et al. (2019). High aluminum stress drives different rhizosphere soil enzyme activities and bacterial community structure between aluminum-tolerant and aluminum-sensitive soybean genotypes. Plant Soil. https://doi.org/10.1007/s11104-019-04089-8

Lugtenberg, B., Kamilova, F. (2009). Plant-Growth-Promoting Rhizobacteria. Annu Rev Microbiol 63, 541–556

Lundberg, D. S., Lebeis, S. L., Paredes, S. H., Yourstone, S., Gehring, J., Malfatti, S., Dangl, J. L. (2012). Defining the core Arabidopsis thaliana root microbiome. Nat. 488(7409), 86–90. doi: 10.1038/nature11237

Lupatini, M., Korthals, G.W., de Hollander, M., Janssens, T.K.S., Kuramae, E.E. (2017). Soil Microbiome Is More Heterogeneous in Organic Than in Conventional Farming System. Front. Microbiol. 7, 2064. doi: 10.3389/fmicb.2016.02064

Maarastawi, S.A., Frindte, K., Linnartz, M., Knief, C. (2018) Crop Rotation and Straw Application Impact Microbial Communities in Italian and Philippine Soils and the Rhizosphere of Zea mays. Front. Microbiol. 9, 1295. doi: 10.3389/fmicb.2018.01295

Magoc, T., and Salzberg, S. L. (2011). FLASH: fast length adjustment of short reads to improve genome assemblies. Bioinformatics 27, 1–8. doi: 10.1093/bioinformatics/btr507

Mendes, R., Garbeva, P., and Raaijmakers, J. M. (2013). The rhizosphere microbiome: significance of plant beneficial, plant pathogenic, and human pathogenic microorganisms. FEMS Microbiol. Rev. 37, 634–663. doi: 10.1111/1574-6976.12028

Moreno-Espíndola, I. P., Ferrara-Guerrero., et al. (2018). The bacterial community structure and microbial activity in a traditional organic milpa farming system under different soil moisture conditions. Front. Microbiol. 9, 2737. doi: 10.3389/fmicb.2018.02737

Mus, F., Crook, M. B., Garcia, K., Garcia Costas, A., Geddes, B. A., Kouri, E. D., et al. (2016). Symbiotic nitrogen fixation and the challenges to its extension to nonlegumes. Appl. Environ. Microbiol. 82, 3698–3710. doi: 10.1128/aem.01055-16

Muzangwa, L., Mnkeni, P.N.S., Chiduza, C. (2017). Assessment of Conservation Agriculture Practices by Smallholder Farmers in the Eastern Cape Province of South Africa. Agron. 7, 46.

Nelson M., Shephard, S., Clifton, M. (2017). The impact of adoption of conservation agriculture on smallholder farmersâ€™ food security in semi-arid zones of southern Africa. Agri. Food Secur. 6, 32. https://doi.org/10.1186/s40066-017-0109-5

Nichols, V., Verhulst, N., Cox, R., and Govaerts, B. (2015). Weed dynamics and conservation agriculture principles: a review. Field Crops Res. 183, 56–68. doi: 10.1016/j.fcr.2015.07.012

Ofek, M., Hadar, Y., Minz, D. (2012). Ecology of Root Colonizing Massilia (Oxalobacteraceae). PLoS ONE 7(7): e40117. https://doi.org/10.1371/journal.pone.0040117

Øyvind H, Harper, David AT, Paul DR. (2001). Past: Paleontological Statistics Software Package for Education and Data Analysis. Palaeontologia Electronica. 4, 4–9.

P.K. Ghosh, A. Das, R. Saha, E. Kharkrang, A.K. Tripathy, G.C. Munda, S.V. Ngachan. (2010). Conservation agriculture towards achieving food security in north east India. Cur. Sci. 99, 915–921.

Parks, D. H., and Beiko, R. G. (2010). Identifying biologically relevant differences between metagenomic communities. Bioinformatics 26, 715–721. doi: 10.1093/bioinformatics/btq041

Quast, C., Pruesse, E., Yilmaz, P., Gerken, J., Schweer, T., Yarza, P., et al. (2013). The SILVA ribosomal RNA gene database project: improved data processing and web-based tools. Nucleic Acids Res. 41, D590–D596. doi: 10.1093/nar/gks1219.

Rahi P, Kapoor R, Young JPW, et al. (2012). A genetic discontinuityin root-nodulating bacteria of cultivated pea in the Indian trans-Himalayas. Mol Ecol. 21(1), 145–59. doi: 10.1111/j.1365-294X.2011.05368.x

Rahi, P. (2017). Mining of Microbial Wealth and MetaGenomics. Phytomicrobiome: A reservoir for sustainable agriculture. Page no 117–132

Rahman, M. M., Islam, A. M., Azirun, S. M., & Boyce, A. N. (2014). Tropical Legume Crop Rotation and Nitrogen Fertilizer Effects on Agronomic and Nitrogen Efficiency of Rice. The Sci World J. 490841. http://doi.org/10.1155/2014/490841

Richardson, A.E., Barea, JM., McNeill, A.M. et al. (2009) Acquisition of phosphorus and nitrogen in the rhizosphere and plant growth promotion by microorganisms. Plant Soil. 321, 305–339. https://doi.org/10.1007/s11104-009-9895-2

Ropitaux, M., Bernard, S., Follet-Gueye, M.L., et al. (2019). Xyloglucan and cellulose form molecular cross-bridges connecting root border cells in pea *(Pisum sativum*). Plant Physiol Biochem. 139, 191–196. doi: 10.1016/j.plaphy.2019.03.023.

Sánchez-Cañizares, C., Jorrín, B., Poole, P. S., and Tkacz, A. (2017). Understanding the holobiont: the interdependence of plants and their microbiome. Curr. Opin. Microbiol. 38, 188–196. doi: 10.1016/j.mib.2017.07.00

Sánchez-Cañizares, C., Jorrín, B., Poole, P. S., and Tkacz, A. (2017). Understanding the holobiont: the interdependence of plants and their microbiome. Curr. Opin. Microbiol. 38, 188–196. doi: 10.1016/j.mib.2017.07.001

Sasse, J., Martinoia, E., and Northen, T. (2017). Feed your friends: do plant exudates shape the root microbiome? Trends Plant Sci. 23, 25–41. doi: 10.1016/j.tplants.2017.09.003

Schöps, R., Goldmann, K., Herz, K., Lentendu, G., Schöning, I., Bruelheide, H., Wubet, T., Buscot, F. (2018). Land-Use Intensity Rather Than Plant Functional Identity Shapes Bacterial and Fungal Rhizosphere Communities. Front. Microbiol. 9, 2711. doi: 10.3389/fmicb.2018.02711

Schreiter, S., Ding, G-C., Heuer, H., Neumann, G., Sandmann, M., Grosch, R., Kropf, S., Smalla, K. (2014). Effect of the soil type on the microbiome in the rhizosphere of field-grown lettuce. Front. Microbiol. 5, 144. doi: 10.3389/fmicb.2014.00144

Suryavanshi, M. V. et al. (2016). Hyperoxaluria leads to dysbiosis and drives selective enrichment of oxalate metabolizing bacterial species in recurrent kidney stone endures. Sci. Rep. 6, 34712; doi: 10.1038/srep34712

Thakuria, D., Hazarika, S., Krishnappa R. (2016) Soil Acidity and Management Options. Indian J Fert. 12, 40–56.

Wagner, M.R., Lundberg, D.S., Del Rio, T.G., Tringe, S.G., Dangl, J.L., Mitchell-Olds, T., 2016. Host genotype and age shape the leaf and root microbiomes of a wild perennial plant. Nat. Commun. 7, 1–15. https://doi.org/10.1038/ncomms12151

Weinert, N., Piceno, Y., Ding, G. C., Meincke, R., Heuer, H., Berg, G., et al. 2011. PhyloChip hybridization uncovered an enormous bacterial diversity in the rhizosphere of different potato cultivars: many common and few cultivar-dependent taxa. FEMS Microbiol. Ecol. 75, 497–506. doi: 10.1111/j.1574-6941.2010.01025.x

Wu, P., Xiong., X, Xu, Z., Lu, C., Cheng, H., Lyu, X. et al. 2016. Bacterial communities in the rhizospheres of three mangrove tree species from Beilun Estuary, China. PLoS ONE. 11(10), e0164082. doi: 10.1371/journal.pone.0164082

Yamamoto, K., Shiwa, Y., Ishige, T., Sakamoto, H., Tanaka, K., Uchino, M., Tanaka, N., Oguri, S., Saitoh, H., Tsushima, S. 2018. Bacterial diversity associated with the rhizosphere and endosphere of two halophytes: *Glaux maritima* and *Salicornia europaea*. Front. Microbiol. 9, 2878. doi: 10.3389/fmicb.2018.02878

Yurgel, S.N., Douglas, G.M., Dusault, A., Percival, D., Langille M. 2018. Dissecting community structure in wild blueberry root and soil microbiome. Front. Microbiol. 9, 1187. doi: 10.3389/fmicb.2018.0118

Zhang, X. M., Wei, H. W., Chen, Q. S., Han, X. G. 2014. The counteractive effects of nitrogen addition and watering on soil bacterial communities in a steppe ecosystem. Soil Biol. Biochem. 72, 26–34. doi: 10.1016/j.soilbio.2014.01.034

Zhernakova, A., Kurilshikov, A., Bonder, M. J., Tigchelaar, E. F., Schirmer, M., Vatanen, T., et al. 2016. Population-based metagenomics analysis reveals markers for gut microbiome composition and diversity. Sci. 352(6285), 565–569. doi: 10.1126/science.aad3369

